# ILLUSORY LIGHT DRIVES PUPIL RESPONSES IN PRIMATES

**DOI:** 10.1101/2024.02.27.582033

**Authors:** Jean-Baptiste Durand, Sarah Marchand, Ilyas Nasres, Bruno Laeng, Vanessa De Castro

## Abstract

In humans, the eye pupils respond to both physical light sensed by the retina and mental representations of light produced by the brain. Notably, our pupils constrict when a visual stimulus is illusorily perceived brighter, even if retinal illumination is constant. Yet, it remains unclear whether such perceptual penetrability of pupil responses is an epiphenomenon unique to humans or whether it represents an adaptive mechanism shared with other animals to anticipate variations in retinal illumination between successive eye fixations. To address this issue, we measured the pupil responses of both humans and macaque monkeys exposed to three chromatic versions (cyan, magenta, and yellow) of the Asahi brightness illusion. We found that the stimuli illusorily perceived brighter or darker trigger differential pupil responses that are very similar in macaques and human participants. Additionally, we show that this phenomenon exhibits an analogous cyan bias in both primate species. Beyond evincing the macaque monkey as a relevant model to study the perceptual penetrability of pupil responses, our results suggest that this phenomenon is tuned to ecological conditions since the exposure to a “bright cyan-bluish sky” may be associated with increased risks of dazzle and retinal damages.

## INTRODUCTION

The eye pupil is the dark aperture in the middle of the iris through which light enters the eye and stimulates the photosensors of the retina. Among vertebrates, pupils exhibit a remarkable variety of circular, elongated, or curved forms, depending on the animals’ specific needs in terms of light exposure, field of view, depth of focus, visual acuity, *etc* (Banks et al., 2015; Malmström & Kröger, 2006). However, they share a common ability of adjusting rapidly to changes in the level of incident light, through the action of the constrictor and dilator iris’ muscles. When incident light becomes scarce, the pupil dilates to increase the surface of light capture and field of view, and when the light intensity rises, it constricts to reduce retinal exposure, prevent dazzle, and produce sharper images and better focus depth. This photomotor response, known as the pupillary light reflex (PLR), is mediated by subcortical structures constituting both a parasympathetic constriction pathway and a sympathetic dilation pathway that have been thoroughly studied in both humans and animals (Diamond, 2001; Mathôt, 2018; McDougal & Gamlin, 2014).

Yet, in humans, accumulating evidence indicates that the PLR cannot be summed up in this simple ‘reflex’ or closed-loop servomechanism driven by the physical light impinging on the retina (Stark, 1962). Besides being modulated by factors such as arousal, attention, motivational states, emotions, or task demands (Ebitz & Moore, 2018; Joshi & Gold, 2020; Peinkhofer et al., 2019), our pupils also respond to mental representations of light constructed by the brain. For instance, when both a bright and a dark visual objects are presented each to a different eye and thus compete for visual awareness (*i.e.* binocular rivalry), the pupils are more constricted (dilated) when the bright (dark) object accesses awareness, despite constant level of stimulating light (Lowe & Ogle, 1966). In the same vein, when the bright and dark objects are presented to both eyes and access visual awareness concomitantly, pupil size reflects which one is covertly attended (Mathôt et al., 2013). In addition, our pupils can also constrict more than expected when exposed to pictures or photos of the sun (Binda et al., 2013), to words such as “bright” (Mathôt et al., 2017), and even in the absence of external stimuli, when we form mental images of bright objects or scenes (Laeng & Sulutvedt, 2013). Finally, brightness visual illusions have also brought very compelling evidence of the perceptual penetrability of the human PLR, since pupil size has been shown to reflect the (illusory) perceived brightness of the stimuli, rather than their physical luminance [*e.g.* (Laeng, Kiambarua, et al., 2018; Laeng & Endestad, 2012)]. This is notably the case with the Asahi brightness illusion created by Akiyoshi Kitaoka, which relies on the phenomenon of optical glare (Tamura et al., 2016). In this phenomenon, the luminance/color gradients of surrounding elements make the center of the image look either brighter (‘glare’ effect) or darker (‘halo’ effect), though the central area is identical in color and luminance in all cases. In its yellow ‘glare’ version (see third line and left column of **Figure 1A**), this illusion has been shown to elicit pupil constriction in humans while they simply look at it, independent of whether they are free to move the eyes or maintain fixation at the center of the pattern or hole (Laeng & Endestad, 2012). Importantly, the amount of pupil constriction was found to correlate with the perceived subjective strength of the glare luminosity as rated by the observers.

**Figure 1.**
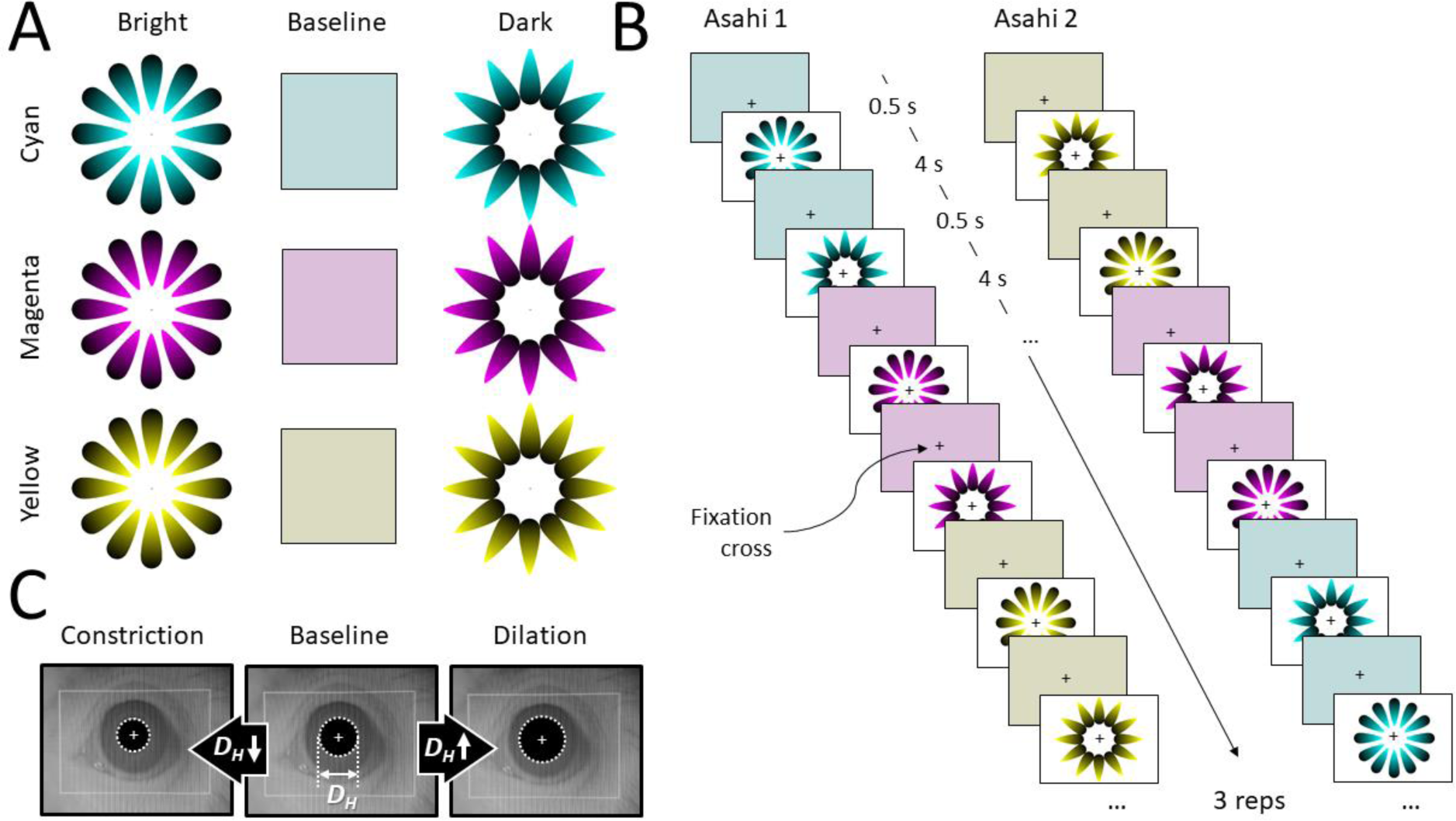
*Experimental task and design*. **A. Stimuli.** The bright and dark versions of the Asahi brightness illusion with their overall mean luminance and color baseline (columns) for the 3 tested colors (cyan, magenta and yellow; rows). **B. Protocol**. Participants maintained their gaze on the central fixation cross while the baseline, bright and dark Asahi stimuli were shown according to one of the 2 sequences shown (Asahi 1 or Asahi 2). The sequence was repeated 3 times in each run for a total duration of about 1 minute and 20 seconds. The Asahi 1 and Asahi 2 runs were interleaved and each participant performed 10 runs of each. **C. Expected pupil responses (illustrated in a monkey eye)**. Relative to the baseline condition, the illusory brighter center of the bright Asahi illusion should lead to pupil constriction (*i.e.* a decrease of its horizontal diameter, D_H_), while the illusory darker center of the dark Asahi illusion should evoke pupil dilation (*i.e.* an increase of D_H_).

Surprisingly, it is still unclear whether this perceptual penetrability is unique to the human PLR or whether it is also present in the PLR of other animals. Another open question is whether this phenomenon plays an adaptative role or whether it is simply an epiphenomenon without any significant function. Recently, it has been proposed that the influence of cognitive/perceptual factors on pupil responses might reflect an anticipation mechanism of probable, upcoming, light stimulation that could guarantee efficient protection against light over-exposure or simply being temporarily dazzled by light, hence suggesting the possibility that this putative predictive response represents an adaptive mechanism that enhances survival in our species (Laeng et al., 2022). In this case, natural selection has likely promoted the same mechanism in a majority of species with movable pupils. It is also likely that this mechanism is color-dependent since being dazzled by the sun is more probable when gazing at a cyan-bluish sky during daylight, than when looking at the magenta and dark-blue colors of the sky at twilight.

Although pupillometry allows testing the effect of brightness illusions in individuals-humans or animals-that cannot use language to report their experience (Laeng et al., 2012), only one study has addressed this question directly, by exposing rats to several versions of the Asahi brightness illusion (Vasilev et al., 2023). Albeit some pupil constrictions were documented in this work, they were moderate both in amplitude and in duration compared to those found in humans, and not systematic across different configurations of sizes and colors. This study also documented a potential neural correlate of this effect in the primary visual (V1) cortex, which was prior to the pupil response and thus consistent with a causal role of this cortical area. Yet, among the two differently colored configurations of the Asahi illusion that evoked clearer pupil responses, only one also evoked differential V1 activation. Besides these mitigated results, a consideration to make is the limitation in using nocturnal animals as an animal model, since rats’ vision is not only as developed as that of diurnal mammals, notably the primates, but they are also exposed to a ‘spectral diet’ (Bosten, 2022; Maule et al., 2023; Webler et al., 2019) or a range of colors that is rather different and more limited in their ecology than that of diurnal species. Moreover, rats do not have a fovea and do not need to move their eyes to focus on an object within the visual field, which might limit the role of perceptual/cognitive information in anticipating changes in the level of illumination. Finally, rats’ ecology typically does not include exposing themselves to direct, high above, sunlight from the sky. Hence, there is a strong need to address this question in animal species that are not only evolutionarily closer to humans but that also share a trichromatic color vision system.

For decades, macaque monkeys have been the most widely used non-human primates to study the visual system (Ostner, 2006), and the symbiosis between human and macaque studies has proven fruitful for gaining a better understanding of its structural and functional underpinnings (Horwitz, 2015). Concerning specifically the pupillary behavior, it has been shown that the PLR is very similar in the two species, making the macaque a good model for unraveling the neural mechanisms that can affect pupil size (Clarke et al., 2003; Pong & Fuchs, 2000). Moreover, it has been demonstrated that the ability of macaques to discriminate brightness and brightness illusions is comparable to that of humans (Huang et al., 2002). Based on the above considerations, in this study, we hypothesized that the Asahi brightness illusion yields pupil constriction in macaques and does so in a manner more comparable to what has been described in humans (Laeng & Endestad, 2012) than in rats (Vasilev et al., 2023). Moreover, another goal and hypothesis of this study, is that the colors of the patterns have differential effects on pupillary responses. Previous studies have revealed that colors have different and systematic effects on pupil responses and, most importantly, that bluish colors yield stronger constrictions than other colors (Thurman et al., 2021). This may reflect an adaptation or adjustment to ecological conditions, where looking directly at the sun in the backdrop of a blue sky is conducive to dazzle and possible damage to the photosensitive substrate. The response to bluish brightness illusions is also strongest (Suzuki, Minami, & Nakauchi, 2019), as one would expect if this illusion, like many others, is a prediction of the next moment (Changizi et al., 2008). Hence, we tested three different colors of the illusory displays (cyan, magenta, and yellow) to investigate the possibility that colors that are prevalently associated in the ecology of both humans and other primates to situations of exposure to glare could yield in the monkeys also the strongest responses to the stimuli colored in cyan, or blue-sky like. We consider that this research could be a first step in paving the way to reveal the neural mechanisms and ecological regularities behind these illusions and the way they trigger pupil responses, throughout the macaque animal model.

## METHODS

### Subjects

#### Informed and naïve human participants

Eight human participants were involved in the present study. Four of them (2 females and 2 males, between 23 and 46 years old) were among the authors of the present study and constituted the informed group since they were aware of its goals. The other four (2 females and 2 males, between 28 and 37 years old) were naïve about the experiment, and they were told that the aim was to study pupil responses to color changes. All participants had normal or corrected-to-normal vision and reported no history of neurological or psychiatric disorders. They provided written informed consent before the experiment, which met the requirement of the ethical principles of the declaration of Helsinki and was approved by the local ethic committee (CLERIT). The experiment was carried out in accordance with the approved guidelines.

#### Monkey subjects

Four female rhesus macaques (M01, M02, M03, and M04) between 8 and 16 years old were also involved in this study. The welfare and care of the animals adhered to the guidelines outlined in the European Union legislation (2010/63/UE) and the French Ministry of Agriculture (décret 2013-118). The project received ethical approval from a local ethics committee and obtained authorization from the French Ministry of Research (MP/03/34/10/09). The macaques were born and raised in captivity. To ensure their well-being, they were housed together in a spacious enriched enclosure, allowing them to engage in natural social interactions and foraging behaviors. Daylight was coming in through opaque-glazed skylights. During feeding times, the macaques were transferred to individual cages. Their diet consisted of standard primate biscuits supplemented with a variety of fruits and vegetables. All four monkeys were equipped with a head-holder implanted on the skull during aseptic surgery, with ceramic screws embedded in the skull (Thomas recordings®) and a cement designed for human bone surgery (Palacos®). The head-holder was used to maintain the animals’ head still with respect to the primate chair during the experimental recordings.

### Experimental set-up

Participants (both humans and monkeys) were installed in an experimental set-up lit only by the stimulation screen that they were facing, with a refresh rate of 60 Hz, a resolution of 1920 x 1080 pixels, and subtending 47 x 31 degrees of visual angle at a viewing distance of 50 cm. Humans were seated in a height-adjustable chair, the head lying on a chin rest to minimize head motion during the experiment. Monkeys were installed in sphinx position within a horizontal primate chair resting on a table. Their head was immobilized during the recordings thanks to the head-holder. We used the eventID software (Okazolab®) to control stimuli presentation. An infrared video-based eye tracker (Iscan®) was used to record the participants’ pupil horizontal diameter as well as horizontal and vertical eye positions monocularly at 120 Hz during stimuli presentation.

### Stimuli

The Asahi stimuli used in the present study are shown in **Figure 1A**. The Asahi pattern of gradients which, somewhat abstractly resembles a pattern of converging plant leaves, was originally designed by Akiyoshi Kitaoka (https://www.ritsumei.ac.jp/~akitaoka/light-e.html), at Ritsumeikan University. The reversed or dark versions were designed by Bruno Laeng by simply rotating 180 degrees each gradient element of the Asahi pattern. In the bright version (leftward column), the center looks brighter than the white background while in the dark version (rightward column), the center appears darker than the white background. Importantly, the bright and dark Asahi stimuli share the same mean luminance and mean color, since they differ only by the opposite arrangement of their components. To serve as a reference for pupil size measurements, we also constructed homogeneous baseline stimuli (middle column), capturing the mean luminance and color of the corresponding Asahi stimuli. The bright, dark and baseline stimuli were produced in three different color versions: cyan (upper row), magenta (middle row), and yellow (lower row) in order to assess also the effect of color on these brightness illusions. The yellow stimulus was selected to match that used in the seminal human pupillometry study (Laeng & Endestad, 2012), while the cyan one was chosen to check for a possible bluish bias in this brightness illusion, as already reported in previous studies (Suzuki, Minami, Laeng, et al., 2019; Thurman et al., 2021). Finally, the magenta was retained based on a previous study with illusory expanding holes since this illusion was found to be the most effective with a magenta background (Laeng et al., 2022). Selecting these 3 colors also permitted to sample the hue space homogenously (with a distance of 120° between all 3 colors in hue space). Thus, in Hue/Saturation/Lightness (HSL) notation, the 9 stimuli shown in **Figure 1A** differ in their average hue (180° for the cyan stimuli, 300° for the magenta ones and 60° for the yellow ones) but they share the same average saturation (28%) and average lightness (80%). Their common luminance on screen was 302 cd/m^2^. All the stimuli also shared the same resolution of 960 x 720 pixels and subtended 20 x 20 degrees of visual angle for the participants.

### Experimental task and design

All the participants performed 1 session of 20 runs, during which they had to maintain their gaze on a black fixation cross (0.6 x 0.6 degrees of visual angle) that was always visible at the center of the stimulation screen. Monkeys received fluid reward for performing the task correctly (*i.e.* for keeping the gaze within ± 2° of the cross center), and we stopped data collection after getting 20 runs in which the monkeys had maintained their gaze in the fixation window for at least 85% of the whole run duration.

Each run contained three blocks in which the six Asahi stimuli (Bright/Dark x Cyan/Magenta/Yellow; see **Figure 1A**) were presented successively for 4 seconds each, always preceded by their corresponding baseline stimulus for 0.5 secs. As illustrated in **Figure 1B**, we alternated runs in which the presentation order was either Bright/Cyan, Dark/Cyan, Bright/Magenta, Dark/Magenta, Bright/Yellow, Dark/Yellow (Asahi 1), or the reversed order (Asahi 2). In total, each run lasted about 1.20 sec, so the entire recording session lasted less than 45 minutes per participant, nevertheless allowing 60 repetitions (20 runs x 3 repetition blocks) for each of the six Asahi stimuli.

### Data processing

For each run, the pupil time course was first detrended using a 2^nd^ order polynomial. Eye blinks, that resulted in a brief loss of eye signal, were discarded together with the 50 ms preceding the blinks (to account for eyelids closure) and the 200 ms following the blinks (to account for eyelids reopening and pupil size stabilization). The discarded signal was interpolated using a shape-preserving piecewise cubic interpolation and the entire time course was then smoothed with a 2^nd^ order Savitzky-Golay smoothing filter (temporal window = 258 ms). For each trial (*i.e.* presentation of an Asahi stimulus), the 4 secs of horizontal pupil diameter raw signal (D_Hraw_) was then extracted and normalized (D_H_) as a function of the median pupil diameter measured in the last 50 ms of the preceding baseline condition (D_H0_) by using the following formula: D_H_ = (D_Hraw_ – D_H0_) / D_H0_ x 100. Thus, D_H_ represents the percent change in pupil diameter relative to that evoked by the immediately preceding baseline stimulus. As illustrated in **Figure 1C**, with pupil constriction relative to baseline, D_H_ will decrease and turn negative. By contrast, with pupil dilation, D_H_ will increase toward positive values.

For each trial, we also computed the horizontal and vertical standard deviation of eye position signals (σ_H_ and σ_V_) to estimate fixation accuracy, and the proportion of interpolated pupil signal due to blinks (PI). Individual trials were excluded from further analyses if at least one of the 3 following conditions was encountered: (1) inaccurate baseline pupil size estimation (*i.e.* D_HO_ more than 3 standard deviations away from the distribution of D_HO_ values evoked by the same baseline stimulus during the session), (2) inaccurate fixation (*i.e.* σ_H_ or σ_V_ more than 3 standard deviations away from the distribution of standard deviations observed across all the trials of the session) or (3) pervasive presence of blinks (*i.e.* more than 15% of the whole trial duration has had to be interpolated due to the presence of blinks).

### Statistics

For each kept trial (3464 out of 4080 in total; 85% of all trials), we computed 2 average values of the horizontal pupil diameter (D_H_), one reflecting the early component of the pupil response, from 0.5 to 2 sec post-stimulus onset, and the other capturing the late component, from 2.5 to 4 sec. For each individual and each of these 2 periods, we assessed whether the pupil diameter was influenced by the brightness illusion, by the stimuli color, and by their possible interaction. This was done by performing 2-way ANOVA tests with the bright versus dark illusion of the stimuli as first factor and their color as second factor. Additionally, we also performed post-hoc 2-sample unpaired t-tests in each individual and for each period to compare the pupil diameter evoked by the bright *versus* dark stimuli separately for the 3 different colors. The same tests were also performed at the group level, by combining all the pupil responses of the human naïve group (n=4), the human informed group (n=4) and the macaque monkey group (n=4). Unpaired 2-sample t-tests were also performed on the bright-dark difference in pupil diameter to compare the strength of the brightness illusion on pupil responses between (1) groups and (2) stimuli color. Finally, we used a 2-way ANOVA to estimate the possible interactions between these factors (*i.e.* groups and color) and one-tailed 2-sample t-tests for post-hoc analyses.

## RESULTS

**Figure 2A** shows how the horizontal pupil diameter (D_H_) evolves over time after the presentation of the Asahi stimuli for one exemplar subject of the human informed group (H1, upper row), one of the human naïve group (H5, middle row) and one of the macaque monkey group (M1, lower row). Responses to the illusory bright and dark stimuli are depicted by solid and dashed lines respectively, in the left, middle and right columns for the cyan, magenta, and yellow stimuli, respectively. Two general observations can be made based on the inspection of these profiles. First, it can be seen that for all types of stimuli, the pupil responses generally start with a transient constriction (a negative dip in the profiles) followed by a dilation before reaching a plateau or at least a slower dilation rate. Second, it appears that pupil responses for the illusory bright and dark stimuli tend to show a systematic deviation, with the bright stimuli profiles (solid lines) consistently below the dark stimuli profiles (dashed lines). This deviation is a clear sign that the pupil response is modulated by the illusory brightness, with a smaller (larger) pupil diameter when the stimulus appears brighter (darker).

**Figure 2.**
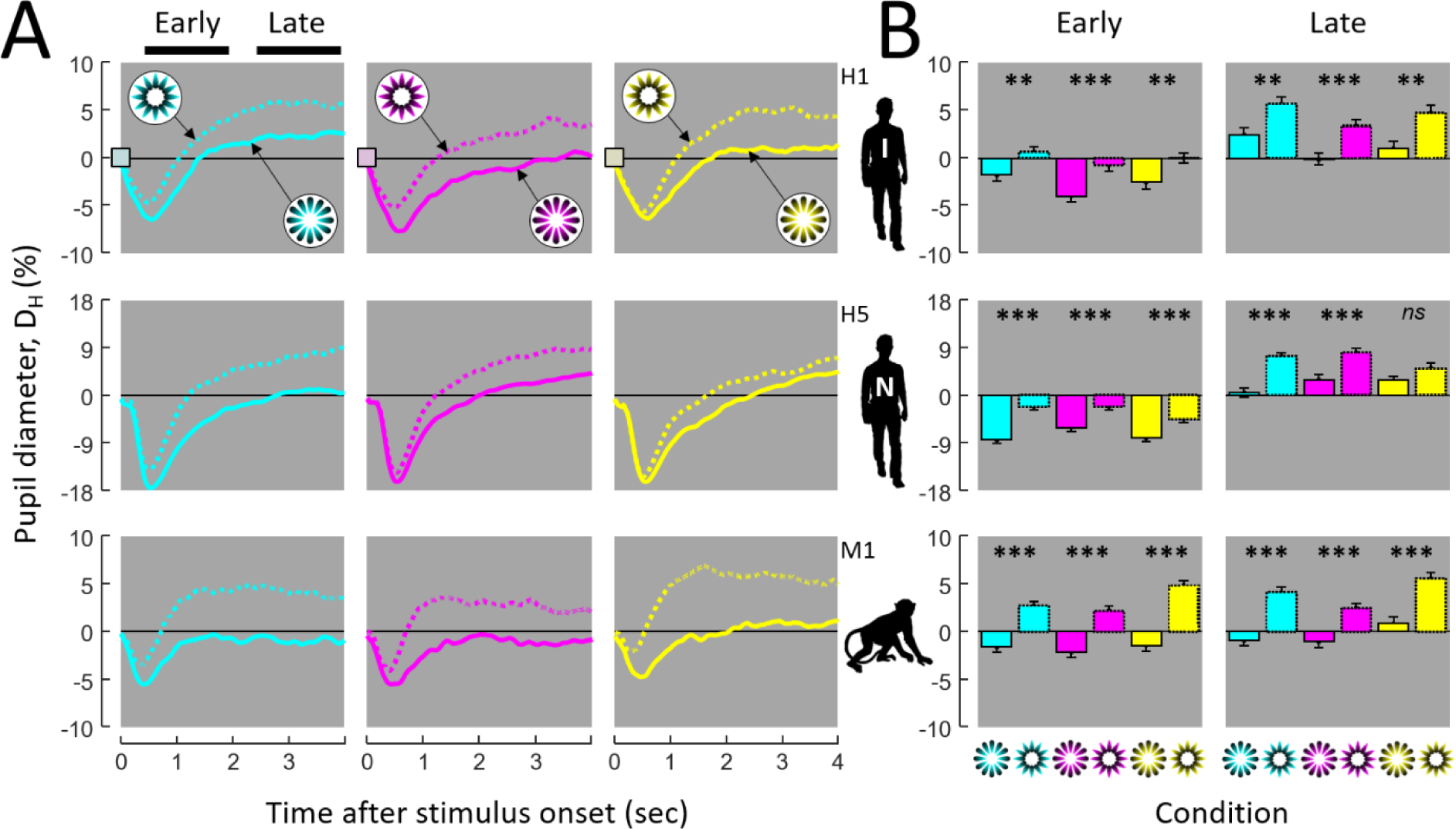
Illustration of individual results. **A.** Mean time courses of the baseline-corrected pupil responses (D_H_) to the bright (solid lines) and dark (dashed lines) Asahi stimuli for 3 exemplar subjects of the human informed and naïve groups (upper and middle rows) and monkey group (lower row). The left, middle, and right columns display the responses to the cyan, magenta, and yellow stimuli respectively. **B.** Mean baseline-corrected pupil diameter integrated over the early (+0.5 to +2.0 sec) and late (+2.5 to +4.0 sec) periods of the responses shown in (A) for the 6 Asahi stimuli. Error bars represent standard errors to the mean over trial repetitions for each subject. Statistical differences between pupil diameters evoked by pairs of illusory bright and dark stimuli were evaluated by unpaired 2-sample t-tests (ns: non-significant, *: p<0.05, **: p<0.01, ***: p<0.001). Human and monkey silhouette images are from PhyloPic.org, available for reuse under Creative Commons licenses. [H: human, M: monkey, I: informed, N: naïve].

From each of these profiles, we computed the mean pupil diameter over an early period (+0.5 to +2 sec) and over a late period (+2.5 to +4 sec) after stimulus onset. These early and late mean D_H_ values are shown in **Figure 2B** and, as expected for these 3 individuals, pupil size was consistently smaller for the illusory brighter stimuli than for their illusory darker counterparts. Unpaired two-sample t-tests (see ‘Methods’) confirmed the statistical significance of this bright versus dark difference in the overwhelming majority of cases. **Figure 3** shows that these observations hold at the group level when averaging results from the 4 individuals constituting each group: informed humans (H1 to H4), naïve humans (H5 to H8), and macaque monkeys (M1 to M4).

**Figure 3.**
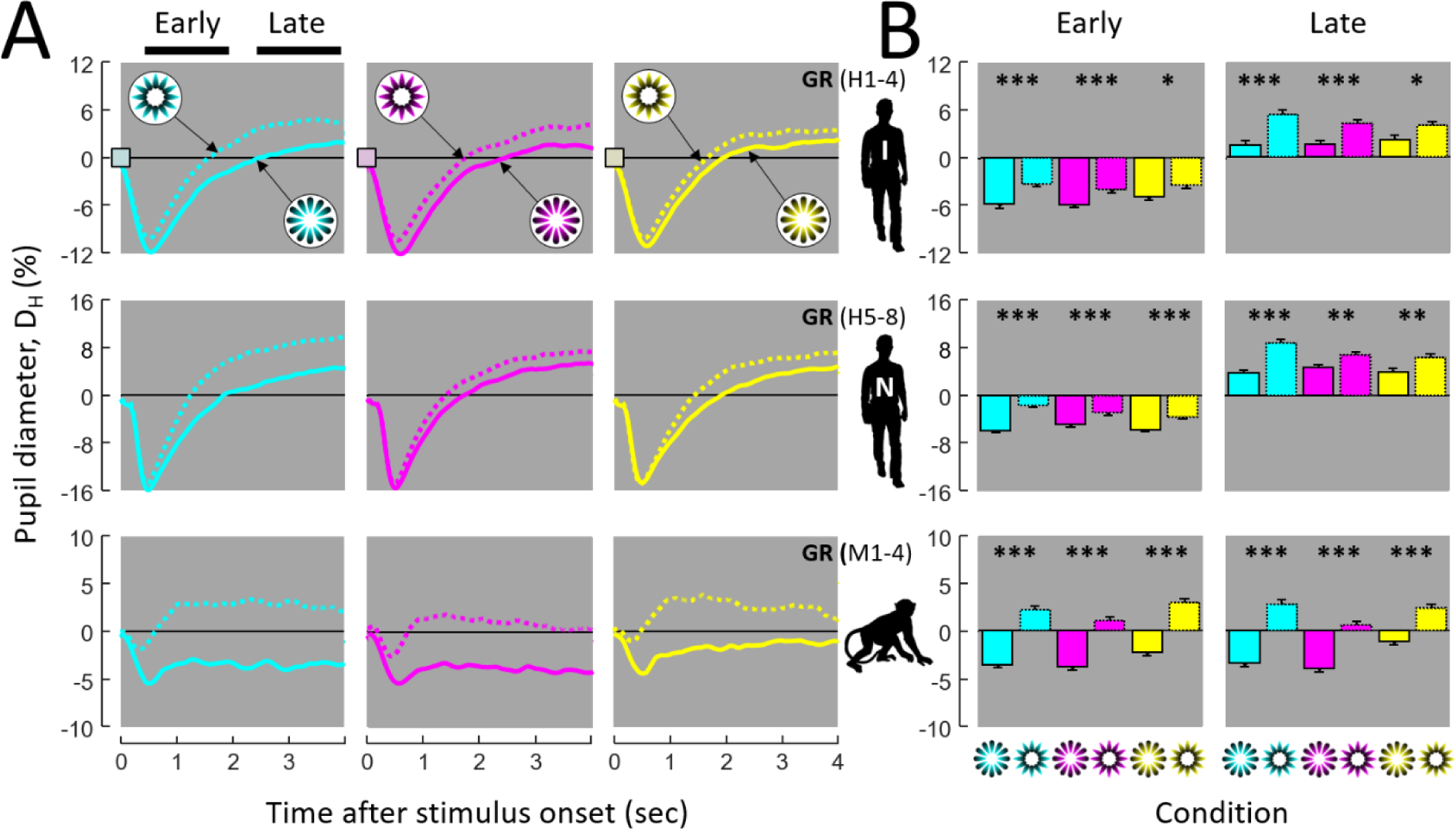
Group results. **A.** Mean time courses of the pupil responses (D_H_). **B.** Mean pupil diameter integrated over the early and late periods of the responses shown in (**A**). Same conventions as Figure 2. [GR: group].

To evaluate whether these observations hold in most individual subjects, we submitted their early and late mean D_H_ values to individual 2-ways ANOVAs, with the illusory bright/dark nature of the stimuli as 1^st^ factor and their cyan/magenta/yellow color as 2^nd^ factor. Results are shown in **Table 1** (columns 1 to 3). A significant effect of the brightness illusion on pupil size was found in 3/4 individuals of the informed human group, 4/4 of the naïve human group, and 4/4 of the macaque monkey group. Unsurprisingly, highly significant results were also found when combining individuals from the same 3 groups (GR in Table 1). Note that the only individual that did not reach significance for the illusion in the 2-way ANOVA (*i.e.* H4) did nevertheless show a significant illusion x color interaction. Performing 2 sample t-tests for each color separately (Table 1, columns 4 to 6 for cyan, magenta, and yellow stimuli respectively) revealed that H4 indeed exhibits a significant effect of the illusion for cyan stimuli in both the early and late phases, but none for both magenta and yellow stimuli. Overall, the 2-sample t-tests shown in **Table 1** indicate that all participants exhibited a significant effect of illusion for at least one color (among the 12 participants x 2 periods, 7/24 cases of 1-color significance, 3/24 cases of 2-colors significance and 14/24 cases of 3-colors significance). Note that at the group level, significance was consistently found for all 3 colors.

**Table 1.** Statistical results of all the individuals and groups (GR). The first 3 columns provide F-values and significance) of the 2-way ANOVA for bright/dark illusion, color and their interaction. The last 3 columns indicate the t-values (and significance) of the 2-sample t-tests between bright and dark stimuli for each of the 3 tested colors (*ns*: non-significant, *: p<0.05, **: p<0.01, ***: p<0.001). [^#^ Subjects represented in Figure 2].

|  |  | 2-way ANOVA |  |  | 2-sample t-test (Bright vs Dark) |  |  |
| --- | --- | --- | --- | --- | --- | --- | --- |
|  |  | Illusion | Color | Inter. | Cyan | Magenta | Yellow |
| **INFORMED** |  |  |  |  |  |  |  |
| H1<sup>#</sup> | E | <u>30.4 ***</u> | <u>4.5 *</u> | 0.3 *ns* | <u>-2.9 **</u> | <u>-3.9 ***</u> | <u>-2.8 **</u> |
|  | L | <u>35.0 ***</u> | <u>5.8 **</u> | 0.1 *ns* | <u>-3.2 **</u> | <u>-3.8 ***</u> | <u>-3.3 **</u> |
| H2 | E | <u>10.7 **</u> | 2.4 *ns* | 0.2 *ns* | <u>-2.4 *</u> | <u>-2.0 *</u> | -1.3 *ns* |
|  | L | <u>11.1 **</u> | <u>4.1 *</u> | 1.5 *ns* | <u>-2.9 **</u> | <u>-2.7 **</u> | -0.4 *ns* |
| H3 | E | <u>8.6 **</u> | 0.9 *ns* | 2.0 *ns* | -0.7 *ns* | <u>-3.5 ***</u> | -1.0 *ns* |
|  | L | <u>16.9 ***</u> | 0.9 *ns* | 0.1 *ns* | <u>-2.0 *</u> | <u>-2.6 **</u> | <u>-2.5 *</u> |
| H4 | E | 3.6 *ns* | 2.1 *ns* | <u>3.7 *</u> | <u>-3.0 **</u> | 0.3 *ns* | -0.4 *ns* |
|  | L | 1.6 *ns* | <u>3.6 *</u> | <u>4.2 *</u> | <u>-2.9 **</u> | 0.1 *ns* | 0.7 *ns* |
| **GR** | E | <u>30.3 ***</u> | **1.4 *ns*** | **0.7 *ns*** | <u>-3.7 ***</u> | <u>-3.4 ***</u> | <u>-2.4 *</u> |
|  | L | <u>44.7 ***</u> | **0.6 *ns*** | **2.2 *ns*** | <u>-5.2 ****</u> | <u>-4.1 ***</u> | <u>-2.3 *</u> |
| **NAIVE** |  |  |  |  |  |  |  |
| H5<sup>#</sup> | E | <u>69.6 ***</u> | <u>5.0 **</u> | 2.2 *ns* | <u>-7.0 ***</u> | <u>-4.0 ***</u> | <u>-3.7 ***</u> |
|  | L | <u>48.6 ***</u> | 2.4 *ns* | <u>3.7 *</u> | <u>-5.9 ***</u> | <u>-4.3 ***</u> | -1.9 *ns* |
| H6 | E | <u>24.3 ***</u> | 0.6 *ns* | 0.5 *ns* | <u>-4.1 ***</u> | <u>-2.7 **</u> | <u>-2.0 *</u> |
|  | L | <u>47.5 ***</u> | 0.2 *ns* | 1.5 *ns* | <u>-5.2 ***</u> | <u>-3.2 **</u> | <u>-3.5 ***</u> |
| H7 | E | <u>9.9 **</u> | 0.9 *ns* | <u>3.3 *</u> | <u>-4.1 ***</u> | -1.1 *ns* | -0.4 *ns* |
|  | L | <u>4.7 *</u> | <u>3.6 *</u> | 1.7 *ns* | <u>-2.7 **</u> | -0.6 *ns* | -0.4 *ns* |
| H8 | E | <u>7.6 **</u> | 0.5 *ns* | 2.7 *ns* | -1.4 *ns* | 0.0 *ns* | <u>-3.5 ***</u> |
|  | L | <u>6.0 *</u> | <u>4.9 **</u> | <u>3.0 *</u> | <u>-2.8 **</u> | 0.6 *ns* | -1.9 *ns* |
| **GR** | E | <u>72.3 ***</u> | **2.9 n.s.** | **4.5 \*** | <u>-7.7 ***</u> | <u>-3.4 ***</u> | <u>-3.8 ***</u> |
|  | L | <u>52.2 ***</u> | **2.0 n.s.** | <u>4.9 **</u> | <u>-6.4 ***</u> | <u>-2.9 **</u> | <u>-3.0 **</u> |
| **INFORMED + NAIVE** |  |  |  |  |  |  |  |
| **GR** | E | <u>95.3 ***</u> | **0.5 n.s.** | <u>4.1 *</u> | <u>-7.7 ***</u> | <u>-4.8 ***</u> | <u>-4.4 ***</u> |
|  | L | <u>88.9 ***</u> | **1.9 n.s.** | <u>6.2 **</u> | <u>-7.9 ***</u> | <u>-4.5 ***</u> | <u>-3.8 ***</u> |
| **MONKEY** |  |  |  |  |  |  |  |
| M1<sup>#</sup> | E | <u>126.8 ***</u> | <u>4.8 **</u> | 2.1 *ns* | <u>-6.4 ***</u> | <u>-5.7 ***</u> | <u>-7.4 ***</u> |
|  | L | <u>83.8 ***</u> | <u>9.1 ***</u> | 0.9 *ns* | <u>-6.4 ***</u> | <u>-4.4 ***</u> | <u>-5.1 ***</u> |
| M2 | E | <u>86.1 ***</u> | <u>5.1 **</u> | 1.0 *ns* | <u>-5.1 ***</u> | <u>-6.0 ***</u> | <u>-5.2 ***</u> |
|  | L | <u>104.9 ***</u> | <u>7.3 ***</u> | 2.1 *ns* | <u>-7.0 ***</u> | <u>-6.9 ***</u> | <u>-4.2 ***</u> |
| M3 | E | <u>171.1 ***</u> | <u>4.3 *</u> | 1.4 *ns* | <u>-7.6 ***</u> | <u>-6.5 ***</u> | <u>-8.8 ***</u> |
|  | L | <u>91.0 ***</u> | <u>4.3 *</u> | 2.3 *ns* | <u>-5.8 ***</u> | <u>-4.7 ***</u> | <u>-6.2 ***</u> |
| M4 | E | <u>45.0 ***</u> | <u>7.3 ***</u> | 1.2 *ns* | <u>-5.0 ***</u> | <u>-3.2 **</u> | <u>-3.3 **</u> |
|  | L | <u>20.5 ***</u> | <u>8.0 ***</u> | <u>4.2 *</u> | <u>-4.4 ***</u> | <u>-2.9 **</u> | -0.5 *ns* |
| **GR** | E | <u>330.8 ***</u> | <u>11.8 ***</u> | **0.8 *ns*** | <u>-11.6 ***</u> | <u>-9.6 ***</u> | <u>-10.5 ***</u> |
|  | L | <u>192.9 ***</u> | <u>15.6 ***</u> | <u>5.2 **</u> | <u>-10.6 ***</u> | <u>-8.0 ***</u> | <u>-5.6 ***</u> |

To get a more general overview, the individuals’ early versus late differences in mean pupil size between the bright and dark illusions for the different stimuli color are shown in **Figures 4A** (informed group), **4B** (naïve group) and **4C** (monkey group). As a first general observation confirming the statistics exposed in **Table 1**, it is clear that for all groups the overwhelming majority of data points lie in the lower left quadrant, meaning that pupil size is smaller for bright illusion stimuli than for the dark ones, in both the early and late periods of the pupil response. Second, for all groups, data points tend to be distributed along a diagonal, which reflects the high degree of correlation between pupil size differences in the early and late periods. Correlation coefficients (r) between the early and late responses for the informed, naïve and monkey groups were 0.76, 0.83 and 0.75 respectively (p < 0.01 in all 3 groups).

**Figure 4.**
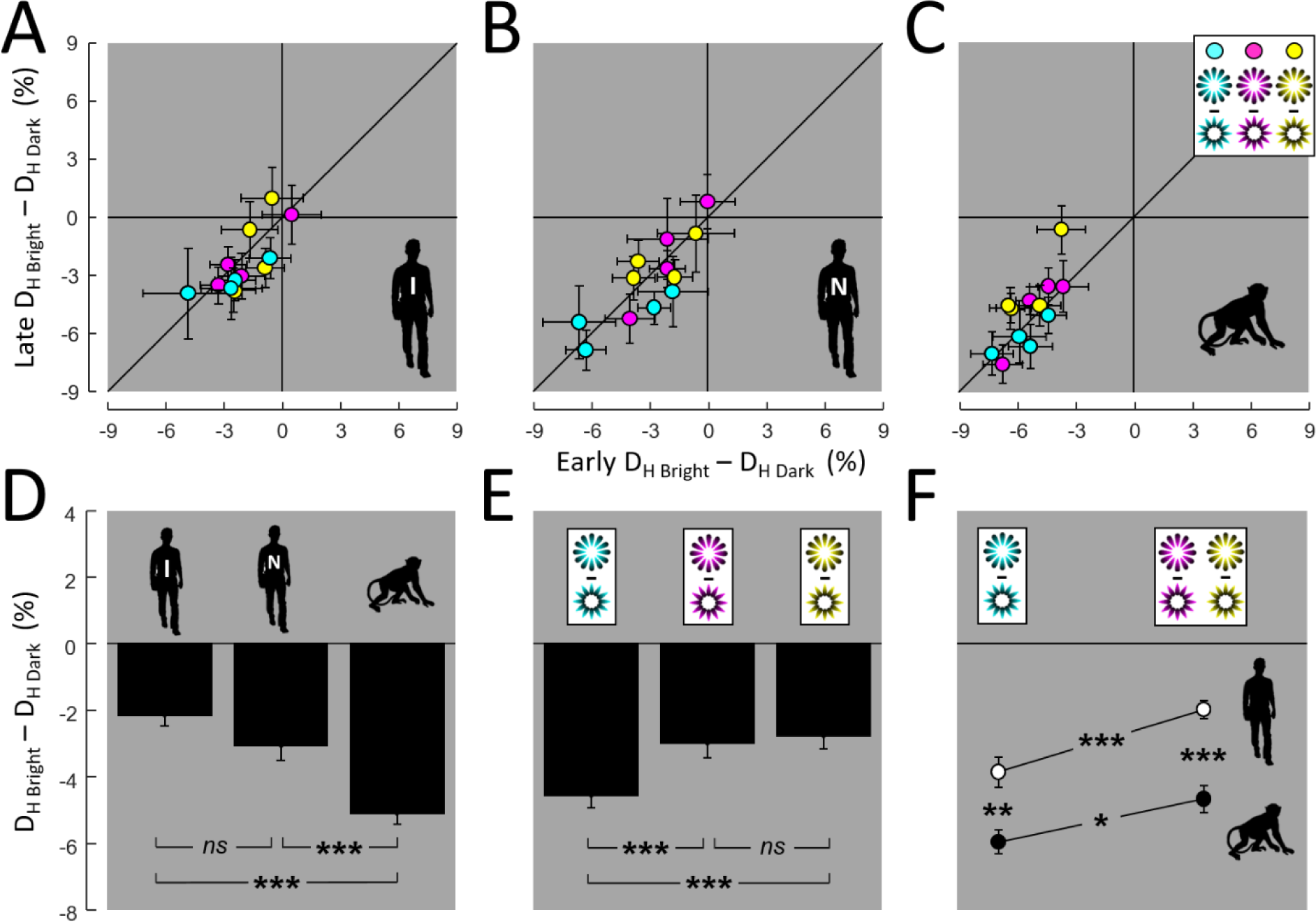
**A-C.** Mean differences in pupil diameter (D_H_) between the bright and dark Asahi stimuli during the early *versus* late periods for all the individuals of the informed human group (**A**), naïve human group (**B**) and monkey group (**C**). Each subject is represented 3 times for the 3 bright/dark pairs in cyan, magenta, and yellow (circular symbols of the same colors, with error bars representing the 90% confidence interval of the mean). **D.** Overall difference in bright-dark pupil diameter for the 3 groups across early and late components and across colors (statistical comparisons with unpaired 2-tailed t-tests). **E.** Overall difference in bright-dark pupil diameter for the 3 color pairs across early and late components and across groups (statistical comparisons with unpaired 2-tailed t-tests). **F.** Overall difference in bright-dark pupil diameter for the cyan versus magenta/yellow stimuli in both humans (informed and naïve subjects together; white symbols) and monkeys (black symbols) subjects (statistical comparisons with unpaired 1-tailed t-tests, with the hypotheses: cyan > magenta/yellow and monkeys > humans). In Figures 4D-F, error bars represent the standard error of the mean. (*ns*: non-significant, *: p<0.05, **: p<0.01, ***: p<0.001).

A closer inspection of **Figures 4A-C** also raises two important observations regarding the effect of the brightness illusion on pupil responses: first, that it might be stronger in monkeys than in both informed and naïve human participants and, second, that it might also be stronger for cyan stimuli than for magenta and yellow ones.

To assess potential group effects, linked to the awareness of the aim of the study (informed versus naïve groups) or to species (human groups versus monkey group), we pooled responses over early and late periods and over colors, as shown in **Figure 4D**. Pairwise comparisons between the groups revealed no significant difference in pupil responses between the informed and naïve human groups (t = 2.04, p = 0.05). By contrast, the difference in pupil size between the bright and dark Asahi illusions was found to be much more pronounced in the monkey group than in both the human-informed group (t = 8.99, p < 0.001) and human-naïve group (t = 4.29, p < 0.001).

We used a similar approach to assess the effect of color by grouping responses over the early and late periods and over the 3 groups, as shown in **Figure 4E**. Pairwise comparisons with 2-sample t-tests indicate that there is no significant difference between the effects observed with the magenta and yellow stimuli (t = −0.61, p = 0.55). However, effects were found to be more pronounced with the cyan stimuli, with respect to both magenta (t = −3.72, p < 0.01) and yellow ones (t = −3.84, p < 0.001).

Finally, we studied whether this cyan preference is expressed differently in humans and monkeys. Since significant difference was found neither between the informed and naïve human groups, nor between the yellow and magenta stimuli, they were pooled to increase statistical power and a 2-way ANOVA was run with species (monkeys versus humans) and colors (cyan vs magenta/yellow) as factors. Results indicate that both factors are highly significant (species: F = 32.5, p < 0.001; colors: F = 14.2, p < 0.001). Importantly, no significant interaction was found between them (F = 0.49, p = 0.49), indicating that pupil reaction to the brightness illusion is stronger in monkeys than in humans and, in both species, for cyan stimuli compared to magenta and yellow ones. These species and color effects are illustrated in **Figure 4F**, with the results of the post-hoc one-tailed t-tests: monkey/cyan > human/cyan (t = 3.0, p<0.01); monkey/(magenta+yellow) > human/(magenta+yellow) (t = 5.6, p < 0.001); monkey/cyan > monkey/(magenta+yellow) (t = 2.0, p < 0.05) and, finally; human/cyan > human/(magenta+yellow) (t = 3.7, p < 0.001).

## DISCUSSION

We compared the pupil responses of humans and macaque monkeys exposed to stimuli of equal luminance but illusorily perceived as brighter (‘glare’ effect) or darker (‘halo’ effect). Our aim was three-fold: (1) to confirm that pupil light responses in humans are not only driven by the intensity of physical light but also by its perceived intensity (i.e., their brightness); (2) to assess whether the perceptual penetrability of pupil light responses is specific to humans or shared with primate relatives and (3) to test whether these brightness effects are color-dependent and in a predictable manner in both human and non-human primates.

Our results firmly establish that pupil responses in humans are strongly modulated by illusory brightness, with significantly greater pupil constrictions for the illusory brighter stimuli than for the illusory darker ones, for all three tested colors. This is in line with the results reported by Laeng and Endestad (Laeng & Endestad, 2012) for the yellow version of the Asahi stimuli as well as with a few other studies (Bombeke et al., 2016; Laeng, Færevaag, et al., 2018). Here, we have extended these findings to other colors (cyan and magenta) and we have shown that the perceptual penetrability of the pupil responses is present in both naïve participants (who thought they were participating in an experiment dealing with color processing) and informed participants (four of the authors of the present study). Crucially, we showed that macaque pupils also respond strongly to illusory brightness. They were found to be even more sensitive to this brightness illusion than humans, showing larger differences in pupil constrictions between the illusory bright and dark stimuli for all three colors. Finally, in both species, we found that cyan stimuli yielded stronger pupil responses to the brightness illusion compared to the yellow and magenta ones. Since the different stimuli versions were equiluminant, one cannot attribute these differences in pupil diameter to changes in the mean luminance of each stimulus. Moreover, because the task performed in this study implied maintaining the gaze on a central fixation cross in both species, all possible explanations related to local luminance in and around the fovea (i.e. eye movements toward different regions of the stimulus) can also be discarded (Clarke et al., 2003; Kardon et al., 1991).

The overall dynamic of the pupil responses to all the Asahi stimuli (see **Figures 2A** and **3A**) is very similar in both species, with a fast constriction during the first half second after stimulus onset, followed by a slower dilation over time throughout the stimulus presentation. The initial pupil constriction is in line with previous studies that proved that the temporal characteristics of the PRL are very similar between both species (Clarke et al., 2003; Douglas, 2018). Although Laeng and Endestad (Laeng & Endestad, 2012) reported faster constriction peaks, this difference could be explained by the modifications we applied to the methodology (*e.g.* color baseline stimuli rather than a gray or blank screen as the baseline inter-stimuli condition). A subsequent study by Suzuki and colleagues (Suzuki, Minami, & Nakauchi, 2019) showed pupil waveforms and peaks more similar to the present results. The subsequent dilation of the pupils can be explained as an effect of the attentional process that the task requires (*e.g.* target fixation) (Beatty, 1982; Laeng et al., 2012; Peinkhofer et al., 2019) and possibly a building up over time of a ‘surprise’ response to the stimuli, which could pull the pupil diameter towards a dilation response (Preuschoff et al., 2011). The pupil seems to reach a plateau earlier in monkeys than in humans, which is also in line with a previous study documenting faster recovery from constriction in macaques (Gamlin et al., 1998).

Importantly, the differences in pupil size between the bright and dark stimuli arise early in these pupil responses and they are maintained with roughly constant strength during the 4 seconds of stimulus exposure (see **Figures 4A-C**). The dynamic of pupil susceptibility to the brightness illusion found here in both humans and macaque monkeys is in line with that reported for humans by Laeng and Endestad (Laeng & Endestad, 2012) (see the lower left time courses in their Figure 3). However, it differs from the dynamic described in rats with comparable Asahi stimuli (Vasilev et al., 2023), since the bright versus dark differences, when present, were mostly expressed in the initial component of the pupil responses.

Although the pupil responses were largely comparable between humans and macaque monkeys, we also found that the overall difference in pupil size between the bright and dark Asahi stimuli was significantly more pronounced in macaque monkeys. In humans, a correlation has been established between the subjective strength of the brightness illusion, as estimated by the participants, and their difference in pupil size for the bright and dark Asahi stimuli (Laeng & Endestad, 2012). Thus, it is tempting to postulate that the monkeys perceive the illusory brightness more vividly than humans do. Nonetheless, we must be cautious with this term in animal studies, given that we have no direct access to the content of the animals’ subjective perception. However, a comparative term could be sensing (Charbonneau et al., 2022), defined as basic information processing (with absent awareness) focused on visual sensory information elicited by the brightness illusions and measured by pupillometry. Although it has been suggested that S-cones might contribute differently to luminance processing in macaques and in humans (Horwitz, 2015), there is no evidence to date supporting the view that macaque monkeys could have a higher sensitivity to (illusory) brightness (Huang et al., 2002). An alternative explanation might be that the PLR differs between both primate species in the sense that monkeys exhibit greater pupil constriction (dilation) in response to an increment (decrement) of incident light intensity or perceived brightness. However, to our knowledge, there is to date no evidence to support this view either (Douglas, 2018; Pong & Fuchs, 2000). Another possibility is that the pupil response to glare is based on the species’ history of exposure to glare; in such a case, one could envision that the classic scenario where the Asahi is a schematic figure of the sun viewed through a canopy of leaves is closer to an experience that is more typical of the ecology of monkeys than of humans. In this sense, the animals tested in the present study were never exposed to experiences of looking at the sky, as in nature, hence it may be that this strong pupillary response is an innate characteristic. Finally, recent work has pointed to the inter-individual variability in pupil size at rest as a possible marker for some cognitive abilities (Aminihajibashi et al., 2019) and there is debate as to whether this factor might predict pupil responses to brightness illusions (Sulutvedt et al., 2021; Wardhani et al., 2022). Thus, we asked whether baseline pupil size for the 3 baseline conditions could predict the differences in pupil susceptibility to the brightness illusion between humans and macaque monkeys. This looked especially interesting since the only human (i.e. H4) who did not have a main effect of illusory brightness, had the largest pupil size during baseline conditions compared to the other human participants. However, we found no correlation between baseline pupil size and pupil susceptibility to the brightness illusion (‘bright’ – ‘dark’ pupil size) over all participants and colors (r = 0.04, p = 0.89). Furthermore, there was no significant difference in baseline pupil size between human participants and macaque monkeys (2-sample t-test, t = 0.12, p = 0.91), in agreement with a recent work (Selezneva et al., 2021). Hence, baseline pupil size can hardly account for any of the differential brightness effects reported here between humans and monkeys.

The other relevant result of the present study is the greater efficiency of the cyan stimuli, compared to the yellow and magenta ones, to elicit pupil responses to illusory brightness. Importantly, this cyan bias was observed in both primate species (see **Figure 4F**), revealing a cross-specific chromatic influence on the brightness illusion (Corney et al., 2009). Note that until this study, humans had been tested only with the yellow version of Asahi (Laeng & Endestad, 2012), though analogous, colored, glare stimuli have been used in other studies and also showed a maximal constriction effect with a blueish color [e.g. (Suzuki, Minami, Laeng, et al., 2019)]. On the other hand, rats were found to be more sensitive to green than to yellow versions of the Asahi illusion (Vasilev et al., 2023). Since no difference was found in baseline pupil size between the cyan, yellow, and magenta baseline stimuli, this factor can also be ruled out as a possible explanation of the cyan bias. This shared cyan bias could also be driven by melanopsin-containing retinal ganglion cells (Gamlin et al., 2007), especially in its late, sustained, component, and is consistent with the fact that color-detection thresholds are largely similar between both species (Gagin et al., 2014).

Considering all the above remarks, the preference observed in both species towards the cyan brightness illusion on pupillary responses in both senses, that is, stronger brightness effect and stronger color effect, leads us to think of an ecological significance, being well preserved across species, at least primate ones. Within this context, our results endorse this ecological approach where the influence of daylight colors, the spectral diet, and/or the “blue sky effect” on pupil constrictions comes from natural experiences where the brain anticipates and prevents possibly being dazzled by the sun (Lafer-Sousa et al., 2012; Thurman et al., 2021). Thus, this strong miosis driven by brightness illusions is important as it can be considered an evolutionary adaptation mechanism protecting the eyes from possible damage by excessive light. In other words, because the perception of light is not straightforwardly related to physical parameters, visual perception may internalize ecological regularities or the natural statistics of the visual world and generate perceptual hypotheses that are most likely to achieve behavioral success (Purves et al., 2004, 2011, 2014). For example, whenever the visual system encounters a geometric pattern of converging gradients, as with the “Asahi” illusion, because it is prepared from individual experience (or that of the species) to the ecological condition of seeing sunlight through cloud formations or plant leaves, it generates a prediction of probable strong source of light in the next moment and such a prediction becomes the perception or best explanation of the current sensory input [see (Brown & Friston, 2012; Changizi et al., 2008; Yon & Frith, 2021)], which leads the eye pupils to adjust adaptively to the percept (Laeng et al., 2022; Zavagno et al., 2017). By doing so, the visual system may respond more promptly and appropriately to potentially risky situations like being temporarily blinded (dazzled) by impinging strong light.

Thus, we have demonstrated that macaque monkeys exhibit pupil responses to illusory brightness. The temporal dynamic and chromatic specificity of these responses fit those we found in humans exposed to the same experimental conditions. These results firmly established the macaque monkeys as a relevant model for studying the neural circuits subtending the perceptual penetrability of the PLR. Moreover, previous monkey studies have established that the perception of brightness illusions correlates with neuronal activity in the early visual cortex (Roe et al., 2005; Salmela & Laurinen, 2007). Further, V1 neurons responses reflect perceived brightness, rather than responding strictly to light (Kinoshita & Komatsu, 2001; Rossi et al., 1996). In the same line, studies with humans show strong activity of V1 and V2 for brightness perception, and not only for luminance (Boyaci et al., 2007; Haynes et al., 2004; Pereverzeva & Murray, 2008). However, some other studies do not support the implication of these early visual areas neither in humans nor in monkeys (Cornelissen et al., 2006; Perna et al., 2005; Ruff et al., 2018), but rather point to the involvement of higher visual areas, like the lateral-occipital sulcus, the intraparietal sulcus, and V4 (Bushnell et al., 2011; Perna et al., 2005; Zhou et al., 2020). Additionally, it has been suggested that brightness perception in both species implies interactions between various cortical regions (prefrontal, frontal, insular, and cingular) and subcortical structures, namely the locus coeruleus and the superior colliculus (Joshi et al., 2016; Peinkhofer et al., 2019). Moreover, V1 and V4 are involved in color coding (Bushnell et al., 2011; Lafer-Sousa et al., 2012; Taylor & Xu, 2022; Yoshioka et al., 1996), and it has been shown that specifically in macaques exist strong V1 responses to orange-cyan, suggesting that V1 is adapted to daylight colors (Lafer-Sousa et al., 2012). What is more, in humans it has been proven that pupil constriction and brightness are already linked to V1 (Bombeke et al., 2016; Suzuki, Minami, & Nakauchi, 2019). Given all the above results, there is still a long way to go to understand all the underlying neural mechanisms.

## CONCLUSIONS

We have shown that our pupils react not only to the intensity of physical light but also to the intensity of perceived light. Importantly, illusory brightness evokes pupil responses that are very similar in humans and macaque monkeys, which notably share a common color-dependency of the effect. Overall, these results suggest that the perceptual penetrability of the pupil light responses is a general, and probably adaptative mechanism, whose underlying neural circuits can now be addressed in a relevant animal model.

## ACKNOWLEDGMENTS

This work was funded by grants from the Centre National de la Recherche Scientifique (CNRS) and from the Agence Nationale de la Recherche (ANR-18-CE37–0022). The authors thank the staff of the Cerco monkey facility (M. Mercier, C. Martinez and A. Leal Rodriguez) for their help with the care and handling of the animals.

